# Where can a place cell put its fields? Let us count the ways

**DOI:** 10.1101/2019.12.19.881458

**Authors:** Man Yi Yim, Lorenzo A Sadun, Ila R Fiete, Thibaud Taillefumier

## Abstract

A hippocampal place cell exhibits multiple firing fields within and across environments. What factors determine the configuration of these fields, and could they be set down in arbitrary locations? We conceptualize place cells as performing evidence combination across many inputs and selecting a threshold to fire. Thus, mathematically they are perceptrons, except that they act on geometrically organized inputs in the form of multiscale periodic grid-cell drive, and external cues. We analytically count which field arrangements a place cell can realize with structured grid inputs, to show that many more place-field arrangements are realizable with grid-like than one-hot coded inputs. However, the arrangements have a rigid structure, defining an underlying response scaffold. We show that the “separating capacity” or spatial range over which all potential field arrangements are realizable equals the rank of the grid-like input matrix, which in turn equals the sum of distinct grid periods, a small fraction of the unique grid-cell coding range. Learning different grid-to-place weights beyond this small range will alter previous arrangements, which could explain the volatility of the place code. However, compared to random inputs over the same range, grid-structured inputs generate larger margins, conferring relative robustness to place fields when grid input weights are fixed.

**Significance statement:** Place cells encode cognitive maps of the world by combining external cues with an internal coordinate scaffold, but our ability to predict basic properties of the code, including where a place cell will exhibit fields without external cues (the scaffold), remains weak. Here we geometrically characterize the place cell scaffold, assuming it is derived from multiperiodic modular grid cell inputs, and provide exact combinatorial results on the space of permitted field arrangements. We show that the modular inputs permit a large number of place field arrangements, with robust fields, but also strongly constrain their geometry and thus predict a structured place scaffold.

## Introduction

Hippocampal place cells exhibit spatially localized firing fields at reproducible locations within and across 2D environments (O’Keefe and Dostrovsky, 1971) and more generally when animals move through more abstract conceptual spaces Aronov et al. (2017). As a population, place cells exhibit complex responses that depend on both environmental context and location: Place cells respond in different spatial combinations in different environments, a process referred to as global remapping Colgin et al. (2008). Within sufficiently large single environments, place cells exhibit multiple fields (Fenton et al., 2008; Park et al., 2011; Rich et al., 2014). What factors determine where place fields will form, and can we predict field locations?

Place cells combine multiple streams of information. In addition to their sensitivity to environmental context, they can also respond strongly to external spatial landmarks and to rewards when such signals are present Hollup et al. (2001); O’Keefe and Burgess (1996). However, when external sensory signals are not present or are spatially uninformative, place cells nevertheless respond reliably and repeatably O’Keefe (1976); Quirk et al. (1990). Interpreting in a neural setting the classic ideas of cognitive maps O’Keefe and Nadel (1978); Tolman (1948); McNaughton et al. (2006) and models of simultaneous localization and mapping (SLAM) Leonard and Durrant-Whyte (1991); Milford et al. (2004); Cadena et al. (2016); Cheung et al. (2012); Widloski and Fiete (2014); Kanitscheider and Fiete (2017c,b,a), we espouse the view that a likely role of place cells is to develop appropriate bindings (maps) between an internal self-consistent scaffold of spatial coordinate representations (presumably generated by grid cells) and external sensory cues (including rewards in the world). In this view, low-dimensional structured grid responses induce complex place cell activity patterns, rather than the alternate view in which simple place field patterns induce grid responses Dordek et al. (2016). The combination of motion-based internal positional estimates with external sensory cues such as landmarks and boundaries into a self-consistent place-cell map enables correction of location estimation errors when the external cues are present Welinder et al. (2008); Burgess (2008); Sreenivasan and Fiete (2011). The same associations further enable intrinsic error correction and location inference in regions where external spatial cues are absent or missing Sreenivasan and Fiete (2011).

Within this perspective, place cells will respond to external cues when they are available, while between encounters with localizing external cues, the dominant reproducible part of their spatially modulated response will be based on self-motion integration, presumably through grid cell activity. In our view, therefore, grid cells induce place cell activity and structure rather than the other way around (cf.While theoretically, place cells could generate internally stable fields (not reliant on grid or external sensory cues) primarily through their own local recurrent dynamics, the capacity of non-modular recurrent networks is extremely small Tsodyks et al. (1996); Samsonovich and McNaughton (1997); Sompolinsky and Kanter (1986); Amit et al. (1985). We believe such mechanisms cannot account for the majority of internally stable place fields, and that most such fields are determined by grid-like inputs. Thus, our focus is on exploring the capacity, constraints and structures of place-field arrangements driven by multi-periodic grid cell inputs, which autonomously generate a large and rich set of input states Fiete et al. (2008); Sreenivasan and Fiete (2011); Mathis et al. (2012) even in the absence of external cues Hafting et al. (2005) through their internal recurrent dynamics Burak and Fiete (2009); Yoon et al. (2013). (At the end, we turn to the contribution of external spatial cues.) Specifically, we consider whether place cells can make free choices in forming stable arrangements of fields given that their inputs are grid-like. At stake is determining whether the underlying scaffold on which place cells associate external cues and location estimates are circumscribed by the geometric nature of the inputs. Mathematically, the problem can be posed as one of determining the capacity of place cells viewed as perceptrons that combine multiple inputs and make a decision on whether to fire (positive label) or not (negative label) by selecting input weights and a firing threshold. Unlike the classical results on perceptrons Cover (1965); Vapnik (1998); Brunel et al. (2004), the inputs in our problem are not random but structured, and not in general position. Being in general position is a property about the linear independence of inputs and is naturally satisfied when these inputs are independently sampled from a smooth distribution. The violation of the assumption that inputs are in general position, which made the counting of dichotomies and the derivation of perceptron capacity tractable Cover (1965), adds complexity to our problem. However, the grid-cell code exhibits a number of symmetries that allow for the computation of some more detailed quantities than typically done for general random patterns, including capacity computations for dichotomies with prescribed numbers of positive labels (fields). Specifically, we will consider ultra-sparse field arrangements, for which the number of fields is fixed, as opposed to merely sparse field arrangements, for which the number of fields scales with some fractional power of the input dimension Itskov and Abbott (2008).

In what follows, we characterize which potential arrangements of place fields are realizable based on grid-like inputs, by providing counting results and deriving stringent theoretical results on the separating capacity or the range of locations over which any arrangement of place fields can be realized. We show that although grid-like inputs are structured, they permit a rich and large repertoire of realizable place-field arrangements over arbitrary ranges. Nevertheless, these arrangements are a small and special subset of potential arrangements and are highly constrained by the input geometry. We show that the separation capacity—over which any potential place-field arrangement is realizable—equals the rank of the grid-like input matrix. We derive an exact rank formula, showing that it approaches the sum of the contributing grid periods. We also favorably compare the stability of place fields with grid-like input relative to place cells with random inputs by examining their optimal separation margins. Our model makes predictions for the relationships between place and grid fields and between adjacent place fields in a place cell. Overall, our results imply that grid-like inputs endow place cells with a large capacity for unique representations, but at the same time place-field configurations are highly constrained by the geometry of grid cells, defining an underlying representational scaffold that can be “overlaid” (that is, tagged or selected) by external cues. Rigorous proofs supporting all our mathematical results are provided in SI Appendix. Portions of this work have appeared previously in conference abstract form (Yim et al., 2019a,b).

## Modeling framework

### Place cells as perceptrons

The perceptron model Rosenblatt (1958) idealizes a single neuron as a linear classifier assigning binary output labels to high-dimensional input patterns 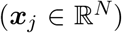 (Fig.1A). The output label is generated by summing entries of the input pattern according to a learned weight vector 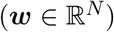 and applying a learned threshold (*θ*):

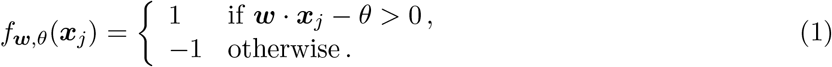

**Figure 1.**
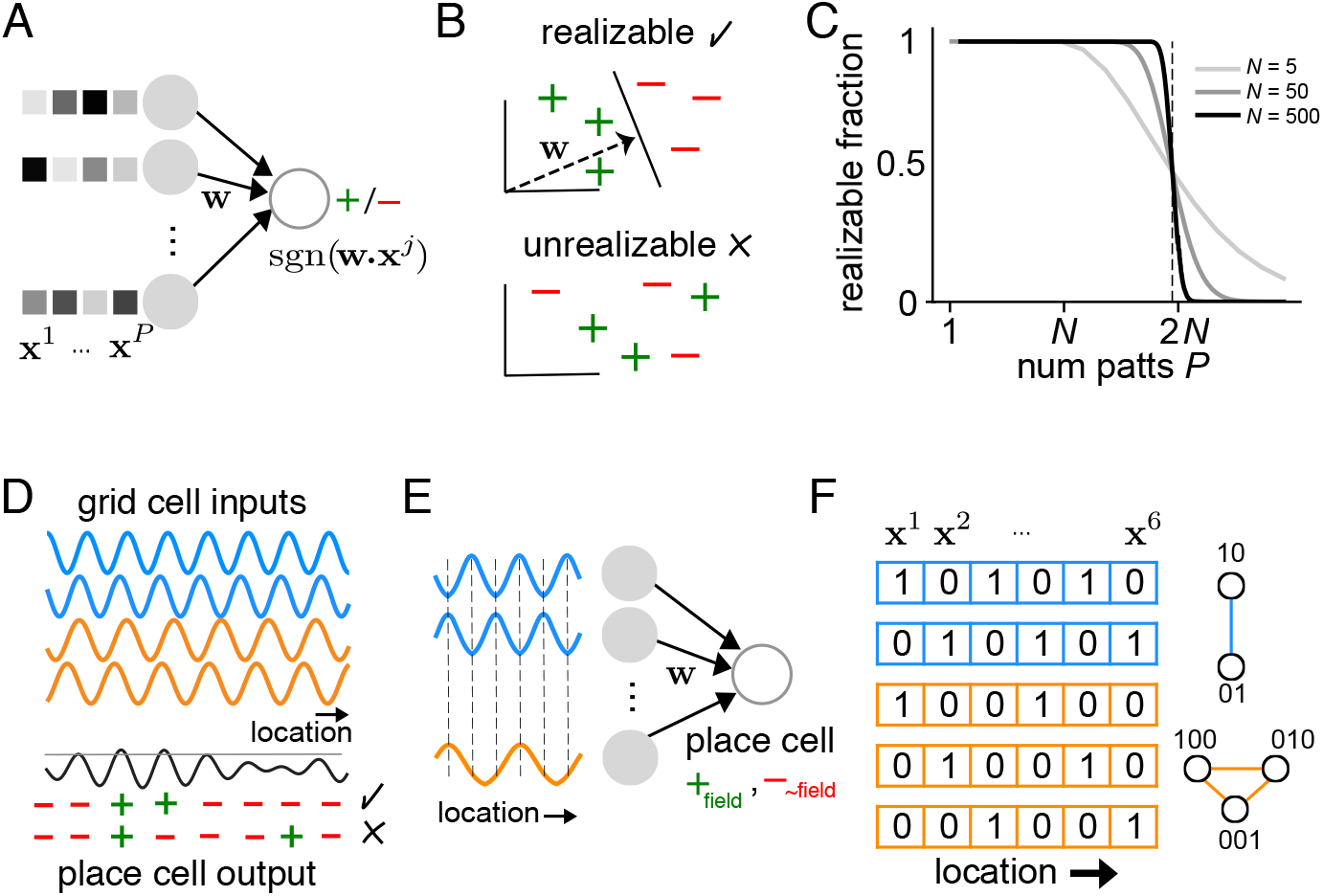
Place cells as perceptrons. (A) A perceptron sums each input pattern (***x**^i^*) with weights *w* and returns a label, +1 or −1. The weights *w* are trained on a set of input patterns and desired labels with the goal that the perceptron labels match the desired labels. (B) A set of patterns can be correctly labeled by a perceptron, and the problem is thus realizable, if the positive examples can be linearly separated (by a hyperplane) from the rest. The perceptron weights *w* encode the direction normal to the separating hyperplane, and the threshold sets its distance from the origin. (C) Cover’s result Cover (1965): For random input patterns, the number of linearly realizable pattern sets divided by the total number of pattern sets as a function of the size of the set is 1 when the set has fewer patterns than the input dimension, and drops rapidly to zero when the number exceeds twice the input dimension; *P* is the number of patterns in the set, and *N* is the input dimension (length of each input pattern). (D) A conceptual view of a place cell as a perceptron: It sums weighted multi-periodic grid cell inputs (blue, orange inputs have different periods; sum shown in black) and can draw a threshold to respond. From this example, it is clear that some field arrangements are realizable (upper row of +, –) while others are not (lower row of +, –). (E) Discretization (in space and response amplitude) of the scenario in (D). (F) Left: Discretized grid-like input code as a function of locations, for two modules with integral periods {2, 3} (blue, orange, respectively). In each module, the number of distinct phases (and cells) equals the period. The total number of locations that can be uniquely represented by the input code (the number of distinct input states or patterns) is the LCM of the grid periods; in this case, 6 (labeled **x**^1^, …, **x**^6^). Right: graphs summarizing the states and state transitions within each module.

A dichotomy is a partitioning of the set of input examples with positive and negative labels. A dichotomy is “realizable” by a perceptron if the positively labeled examples are spatially segregated from the negatively labeled ones by a linear hyperplane (Fig.1B). Cover’s counting theorem provides a count of realizable dichotomies when the patterns are in general position (Cover, 1965). A set of patterns {***x***_1_, …, ***x***_*P*_} in a *N*-dimensional space is in general position if no subset of patterns with less than *N* + 1 patterns is linearly dependent. In other words, no subset of *n* + 1 points lies one a (*n* – 1)-dimensional plane for all *n* ≤ *N*. Cover’s counting theorem states that if the input dimension of the patterns is *N*, then any set of *P* ≤ *S* = *N* + 1 patterns in general position is linearly separable, where *S* is the separating capacity^1^. As *P* grows beyond *S*, the probability that the patterns can be linearly separated decreases, dropping to 1/2 for *P* = 2*N* + 1, Fig. 1C (Cover, 1965).

Here, we consider a place cell as a perceptron whose high-dimensional input patterns (***x**_j_*) are multi-periodic grid-cell patterns for an (appropriately discretized) location *j*. The place cell can make a decision about where to put fields, and equally important, where not to put them, by selecting its weights and threshold: (*f*_***w***,*θ*_(***x**_j_*) = 1: field) or not (*f*_***w**,θ*_(***x**_j_*) = –1: no field), Fig. 1D. A particular *arrangement of fields* across locations is realizable by a place cell if there is some set of input weights and threshold for which the summed grid inputs are above threshold at those locations but below it at all other locations, Fig. 1D. A *K*-field arrangement is any arrangement of exactly *K* fields.

In the following, we ask and answer two distinct but related questions: (1) Out of all potential field arrangements, what fraction is realizable, and how does this realizable fraction differ for grid-like inputs compared to inputs encoded more conventionally, as random or one-hot coded inputs? This is akin to perceptron function counting Cover (1965) with structured rather than general-position inputs. (2) Over what range of locations is every potential field arrangement realizable, and how does this range compare to the range over which the inputs provide a unique encoding of locations? This is analogous to computing the perceptron separating capacity Cover (1965) for structured rather than general-position inputs.

### Structured input patterns

To address these questions analytically, we initially model each grid cell as a {0, 1}-valued periodic function in discretized one-dimensional space, with one of *M* integer-valued periods {λ_1_, …, λ_*M*_}, Fig. 1E. The inputs are arranged in modules, with inputs of one module having a common response period. Cell *i* in module *m* has unit-valued peaks recurring every *ϕ_i_* = (*i* – 1)/λ_*m*_ mod 1, where *i* is a discretized spatial index in one dimension, Fig. 1E. The input module of period λ_*m*_ consists of λ_*m*_ cells with a spread of all possible phases. Thus, each location *j* corresponds to an input pattern ***x**_j_* of {0, 1}-valued vectors of length 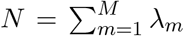, where nonzero entries correspond to co-active grid cells at position *j*. Collectively, these input patterns ***x**_j_* define the *grid-like code*, whose codebook is *X*_g_.

The number of unique input patterns that can be generated by this grid-like code in one spatial dimension is *L* = LCM({λ_1_, …, λ_*M*_}), which grows exponentially with *M* for generic choices of the periods {λ_*m*_} Fiete et al. (2008). We consider field arrangements over the range *l* (1 ≤ *l* ≤ *L*), where the size *L* of the codebook determines the “full range” of the grid-like code. We will consider more realistic models of grid cell activity in later sections.

In the grid-like code, only one cell per module is active at any location. Thus, the grid-like code belongs to a more general class of codes that we call *modular-one-hot* codes. In a modular-one-hot code, inputs are divided into modules; within each module, the code is one-hot (only one active cell per module) and cells are exchangeable. By contrast, cells are not exchangeable across modules if the modules have distinct periods. Modular-one-hot codes are a generalization of grid-like codes in the sense that they exhibit the same modular structure, but there is no notion of a spatial ordering or spatial encoding of locations. Their codebook *X*_mo_ comprises all possible combinations of active cells. These can be written under product form as ***x*** = (***y***_1_, …, ***y**_M_*), where ***y**_m_* is the λ_*m*_-dimensional vector whose only unit component marks the active cell in the *m*-th module. Thus, the codebook *X*_mo_ comprises 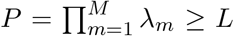 patterns. The codebooks *X*_g_ and *X*_mo_ coincide for pairwise-coprime periods.

Answering question (1) from above involves determining which dichotomies of modular-one-hot patterns can be realized by a perceptron, independent of how these patterns map to spatial locations. Thus, our computations apply generally to modular-one-hot codes and specifically to grid-like codes. The general setting of the modular-one-hot code *X*_mo_, which includes the grid-like patterns *X*_g_, will prove well-suited to perform combinatorial computations.

Answering question (2) from above, which asks about the largest contiguous spatial range over which all spatial field arrangements are realizable, involves determining which realizable dichotomies of modular-one-hot patterns also correspond to realizable dichotomies (arrangements) of grid-like patterns embedded into a given spatial environment. Thus, this question introduces the additional constraint that input patterns represent spatially contiguous positions. In that respect, note that spatially contiguous positions over any range *l* < *L* represent a particular (rather than random) subset of *X*_mo_.

## Geometric arrangements of structured inputs

First, we seek to answer question (1). In modular-one-hot codes *X*_mo_, which generalizes the grid-like code, both the patterns ***x**_j_* (input matrix columns) and the input cells (input matrix rows) are fully exchangeable. This highly symmetric structure determines the geometric arrangement of patterns in *X*_mo_, and therefore also determines the number of dichotomies in *X*_mo_ that are realizable by separating hyperplanes.

To make these ideas concrete, consider a simple example with module periods {2, 3}, Fig 1F (left). In this code, the period-2 inputs alternate between two states: (0, 1) and (1, 0). We represent this activity structure by an edge connecting these two states. Likewise, the period-3 cells alternate between three exchangeable states: (1, 0, 0), (0, 1, 0), and (0, 0, 1), represented by a triangular graph connecting the states, Figs. 1F. Together the modules visit all *P* = 6 combinations of their states; the module updates can be viewed as independent. In other words, *X*_mo_ is geometrically given by the product of the edge graph and the triangular graph, whose convex hull forms an orthogonal triangular prism, Fig. 2A.

**Figure 2.**
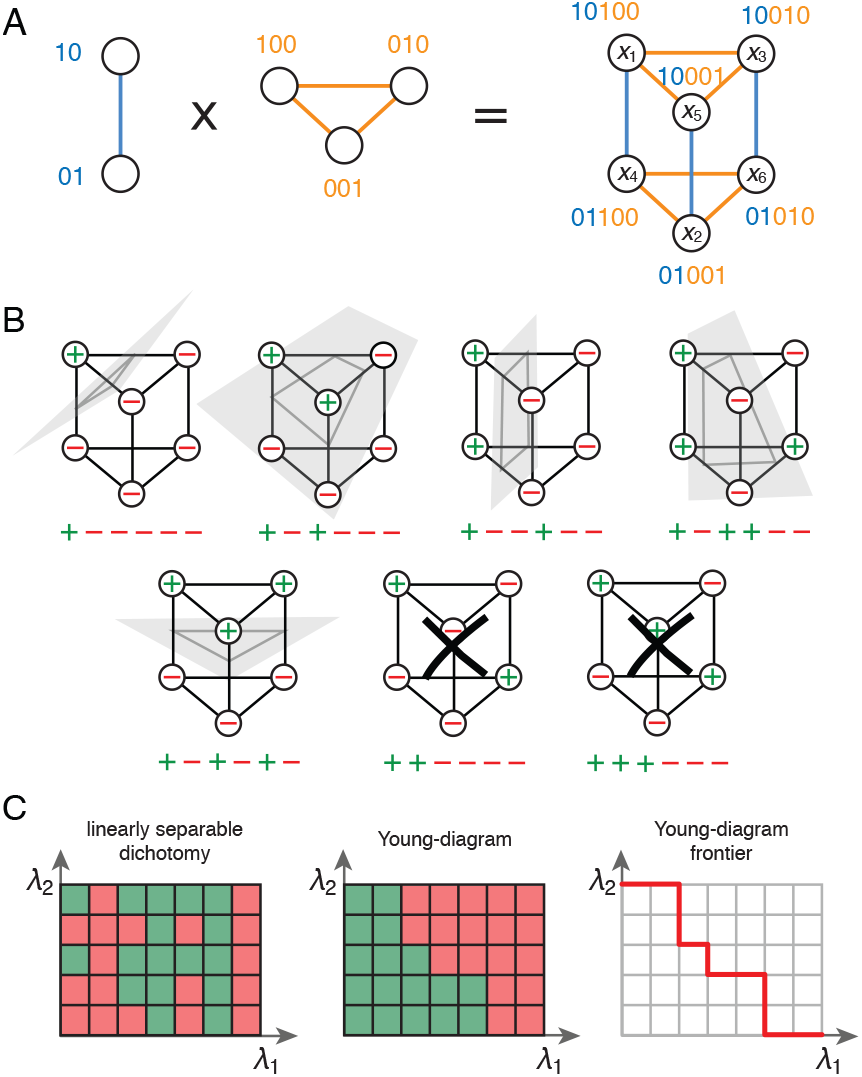
Counting realizable field arrangements in a place-cell perceptron. (A) Geometric representation of grid-like input patterns for the period {2, 3} example of Fig. 1F. Though the input patterns are 5-dimensional, they have a simplified structure given by the product of the 2-graph and 3-graph in Fig. 1F because of the independently-updating modular structure of the code, and can be embedded as a 3D triangular prism. (B) All possible (translationally and symmetrically distinct) labelings of the set of inputs patterns over the full range of the code (*S* = 6), excluding the trivial all ‘- or all ‘+’ (no fields and all fields) combinations. A ‘+’ label on a node of the diagram represents a place field at that input location, and a fully labeled diagram is one particular arrangement of fields and non-fields across all locations. The first five arrangements are realizable, while the last two (with black crosses) are not. (C) For two modules with periods {λ_1_, λ_2_}, each location can be represented as one box on a λ_1_ × λ_2_ grid, whose coordinates are given by the two phases for that location. The set of desired place fields (a desired field arrangement) is then a set of colored boxes on this grid (left). If a colored block of boxes, non-increasing from left to right and top to bottom, can be constructed by swapping rows and columns (middle; this is a Young diagram), the field arrangement is realizable. For two modules, the existence of a Young diagram is necessary and sufficient for realizability. Right: the *frontier* of the Young diagram, used for counting (SI Appendix).

This geometric picture generalizes to an arbitrary number of modules *M* and to arbitrary module periods λ_*m*_, 1 ≤ *m* ≤ *M*. By exchangeability, each pattern in *X*_mo_ is an extreme point of the convex hull. This convex hull defines a polytope which is the orthogonal product of *M* simplices. Each period-λ_*m*_ module is represented by a simplex of dimension λ_*m*_ – 1, 1 ≤ *m* ≤ *M* (SI Appendix). In a convex polytope, any vertex can be separated from all the rest by a hyperplane. Moreover, pairs of vertices can be separated as well, if and only if they are connected by an edge. Thus, geometric insight directly reveals the structure of realizable *K*-dichotomies in *X*_mo_ for *K* = 1 and *K* = 2 (where in a *K*-dichotomy, exactly *K* of the input patterns are positively labelled), showing that all 1-field and 2-field combinations are realizable.

In the following, we further leverage geometric insight to count generic *K*-dichotomies via combinatorial arguments. The many symmetries of the modular-one-hot code will greatly facilitate the enumeration of geometrically distinct dichotomies; for instance, in the period-{2, 3} example, the symmetries reduce 2^6^ = 64 possible dichotomies into only 8 distinct classes (including the all-zeros dichotomy), Fig. 2B.

## Counting realizable dichotomies

For modular-one-hot codes, a separating hyperplane specified by general weights can be equivalently specified by non-negative weights and an appropriate threshold since all inputs have the same L1-norm (SI Appendix; see Amit et al. (1989); Brunel et al. (2004) for similar result with random inputs). This observation allows for a helpful analogy. The construction of a separating hyperplane with non-negative weights for modular-one-hot inputs can be thought of as the composition of a multi-course meal from a menu of options, with a total price constraint: Each module is one course of the meal, each phase (cell) in a module is a choice of dish for that course, and the weight from that cell is the price of that dish. The goal is to construct a meal by picking one item for the first, second, third, etc. courses, so that the total cost remains at or below the fixed budget (the threshold). The number of topologically distinct separating hyperplanes equals the number of meals that fit the budget. For *M* = 2 modules, one can count the number of meals or separating hyperplanes with the help of Young diagrams, which are commonly used to enumerate partitions of the integers, Fig. 2C (SI Appendix) Fulton and Fulton (1997). Young diagrams allow us to convert the geometric problem of hyperplanes, which can vary infinitesimally but for which only certain variations—those that change the classification of a data point—matter, into an appropriately topological one that involves mere counting.

Using the Young-diagram approach, in the case where place cells receive inputs from two grid modules with periods λ_1_ and λ_2_, the number of realizable dichotomies (for all sparseness *K*) in the full range λ_1_λ_2_ of grid patterns is

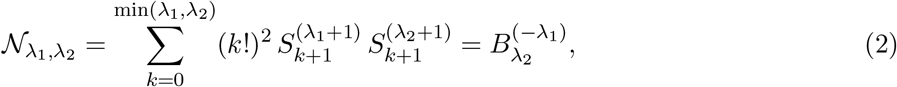

where 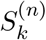 are Stirling numbers of the second kind and 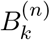 are the poly-Bernoulli numbers (SI Appendix). Poly-Bernoulli numbers were originally introduced by Kaneko to enumerate the set of binary *k*-by-*n* matrices that are uniquely reconstructible from their row and column sums Kaneko (1997). The use of Poly-Bernoulli numbers in enumerating permutations of Young diagrams was pioneered in Postnikov (2006), while their asymptotic behavior along the diagonal was obtained in de Andrade et al. (2015) as:

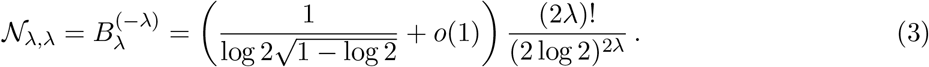

This scales slower than λ^2λ^, while the number of dichotomies scales as 2^λ^2^^. Thus the number of realizable arrangements over the full range is a vanishing fraction of all potential arrangements.

For *M* ≥ 3 modules, it is not possible to directly use Young diagrams: All realizable field arrangements still correspond to Young diagrams, but not all diagrams correspond to a linearly separating hyperplane, thus counting Young diagrams is not equivalent to counting realizable field arrangements. As expected given the difficulty of counting the number of linearly separable Boolean functions of arbitrary (input) dimension, it becomes more difficult to enumerate realizable gridinput-driven field arrangements with arbitrarily many modules.

However, given that place fields are sparse even on long tracks, it is of substantial interest to count the *K*-dichotomies, where *K* is small (*K* = 1, 2, 3 and 4). Using a similar approach, we can count the number 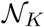 of these small-*K* dichotomies for any number of grid modules in full range, and find that it scales as (derivation of exact number given in SI Appendix):

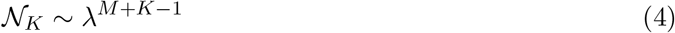

in the limit of large periods λ_1_, …, λ_*M*_ ~ λ → ∞. This is to be compared with the corresponding total number of *K*-dichotomies, which scales as λ^*MK*^. Thus, the fraction of achievable *K*-dichotomies scales as λ^−(*M*–1)(*K*–1)^, which vanishes as a power law for large λ as soon as *M* > 1.

In summary, the set of realizable field arrangements is a vanishingly small fraction of all field arrangements, meaning that place cells, when well-modeled as perceptron readouts of grid cell inputs, have little freedom in where to place their fields.

### Comparison with other input patterns

To give context to our results with grid-like patterns, we introduce some key alternative structured encoding patterns for comparison: How does the number of realizable field arrangements depend on the structure of these patterns, and their level of modularity/hierarchy?

We define the *N*-dimensional *one-hot* (or ‘grandmother-cell’) code, denoted by *X*_oh_, as patterns of input dimension *N* in which there is exactly one active cell, and each cell is active in at most one pattern; and the *N*-dimensional *binary* code, denoted by *X*_b_, where the patterns correspond to Number and fraction of linearly separable dichotomies for the binary, modular-one-hot (*M* = 2), and one-hot codes with the same cell budget.

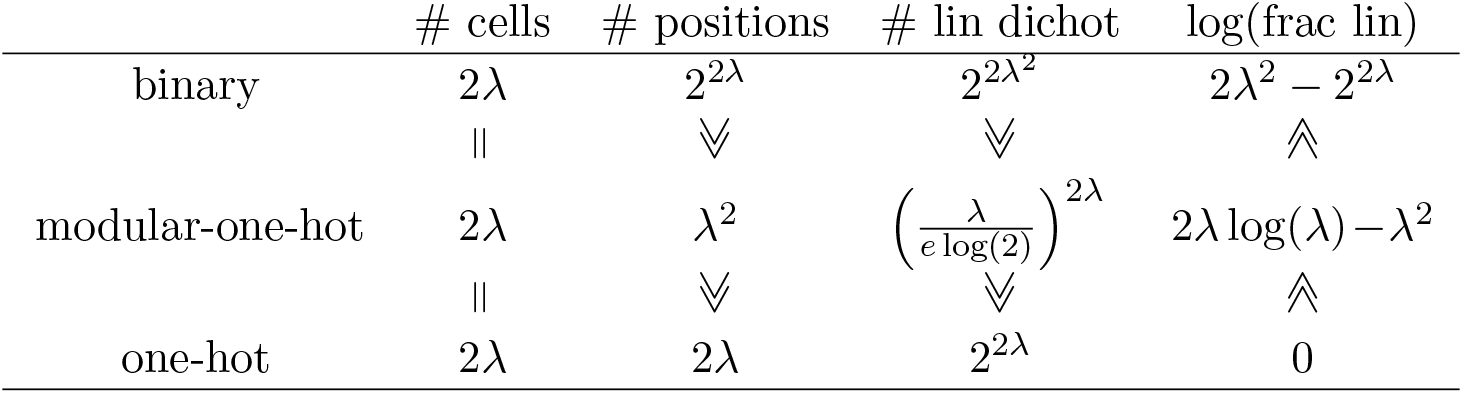

all possible binary activity vectors of length *N*:

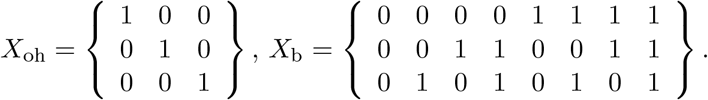

In the one-hot code, all patterns lie on the vertices of a simplex, thereby in general position. All cells are completely exchangeable, and the code has no modularity or hierarchy. It may be viewed as a binarized version of the random patterns normally used for perceptrons: The set of *N* random input patterns is linearly independent, below the separating capacity of *N* + 1.

The binary code is strongly modular and hierarchical: Each cell represents a specific position (register) in the binary number system. Although input patterns are still exchangeable, there is no longer exchangeability of cells as they each represent specific size scales; each cell is its own module. Patterns in the binary code lie on the vertices of a hypercube and thus contain square faces, prototypical examples of a non-general configuration, even for small numbers of patterns relative to number of cells.

The modular-one-hot code, including the grid-like code, exhibits an intermediate degree of hierarchy. Patterns lie on the vertices of a product of simplices; the resulting polytopes with *M* modules contain *M*-dimensional hypercubes and are thus not in general position.

In Table 1, we directly compare the number of realizable dichotomies across codes, while keeping the cell budget, i.e. the input dimension, fixed across codes. Although we expect our comparison to remain valid for an arbitrary number of modules *M*, we have restricted our analysis to the case *M* = 2, for which we have an explicit formula counting the total number of realizable dichotomies. We set λ_1_ = λ_2_ = λ in the modular-one-hot code for ease of direct comparison with the one-hot and binary codes of equal input dimension *N* = 2λ. For large λ, the one-hot code comprises far fewer patterns (*N* = 2λ) than the grid-like code, which in turn comprises far fewer patterns than binary codes (*N*^2^/4 = λ^2^ and 2^*N*^ = 2^2λ^, respectively), Table 1 (second column). This is due to the greater expressive power permitted by modular (grid-like or modular-one-hot) and hierarchical (binary) codes.

How many of these patterns admit realizable dichotomies? Just as for the modular-one-hot codes, the patterns of the one-hot and binary codes fall on the vertices of a convex polytope. For the one-hot code *X*_oh_, the convex hull of patterns is the canonical (*N* – 1)-dimensional simplex, thus any subset of *K* vertices (1 ≤ *K* ≤ *N*) specifies a (*K* – 1)-dimensional face of the simplex and is a linearly separable dichotomy. All 2^λ*M*^ dichotomies of the one-hot code are therefore linearly separable, as expected from Cover’s counting theorem, Table 1 (third column). And, the fraction of realizable dichotomies for the one-hot code is therefore 1, Table 1 (fourth column).

For the binary code *X*_b_, the convex hull of the 2^*N*^ patterns defines a *N*-dimensional hypercube, thus some dichotomies are not linearly separable (e.g., the pair {(0, 0, 0), (1, 1, 1)}). Counting the number of linearly separable dichotomies on the hypercube vertices, also referred to as linear Boolean functions, has attracted much interest Peled and Simeone (1985); Hegedus and Megiddo (1996). It is an NP-hard combinatorial problem. In the limit of large dimension (*N* → ∞), the number of linearly separable Boolean functions scales as 2^*N*^2^/2^ Zuev (1989), much larger than for one-hot codes (Table 1, column three). However, this number is still a vanishing fraction of all 2^2^*N*^^ hypercube dichotomies, Table 1 (fourth column).

For the modular-one-hot code with *M* = 2, we determined earlier the total number of linear dichotomies as the poly-Bernoulli number 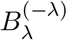. For large periods, the scaling given in (3) is approximately λ^2λ^, permitting a direct comparison with the one-hot and binary codes (Table 1, row 2).

Given a fixed budget of grid cells, a greater degree of hierarchy (modular-one-hot and binary) yields many more potential input patterns, as well as a much larger total number of realizable arrangements (realizable dichotomies), relative to non-modular codes including the one-hot code. At the same time, these realizable dichotomies represent a shrinking fraction of total possible dichotomies, Table 1. The implication is that more hierarchical codes support many more realizable field arrangements, but these are an increasingly special subset of all potential arrangements that are strongly structured by the inputs, and thus cannot be chosen to have random or arbitrary configurations. In summary, when compared with one-hot and binary codes, modular-one-hot codes, including grid-like codes, occupy a middle-ground between constrained structure and pattern richness (permitting many place-field arrangements).

Can we further compare grid-like codes with random (real-valued) codes, for which patterns are always in general position? Such a comparison is hindered by the fact that, in principle, random code can uniquely encode an infinite number of positions, yielding an infinite set of potential dichotomies. To address this caveat, we compare grid-like codes and random codes for the same number of input patterns rather than for the same budget of cells. Then, we seek the required number of cells for the number of dichotomies realizable by the random code to scale as in the grid-like code. Consider *M* = 2 grid modules with large periods of order λ. This corresponds to a range of *P* = λ^2^ input patterns, leading to consider a random code with λ^2^ inputs with a yet-to-be-determined number of cells (input dimension) *N*. As *N* can be assumed small compared to λ^2^, then by the asymptotic form of Cover’s counting theorem, the number of realizable dichotomies scales as *P^N^* = λ^2*N*^ for the random code. Thus, achieving the same scaling as the grid-like code, requires a minimum number of cells of λ. This shows that for *M* = 2, the grid-like code requires a comparable number of cells (twice) to achieve the same number of field arrangements as with random codes. (We conjecture that grid-like codes with *M* modules would require *M* times more cells than random codes over an identical range.)

## Place-cell separating capacity

Above, we considered how many field arrangements a place cell can express when reading out a modular-one-hot code, which remains valid for grid codes when considered over the full range of positions *L*. We now turn to question (2) from above: If we require that all field arrangements be realizable over some contiguous range of inputs, what is the maximum achievable range? This quantity, denoted by *l*\*, is the analogue of Cover’s separating capacity Cover (1965) for grid-like inputs. We therefore call *l*\* the contiguous separating capacity of a place cell.

We provide three primary results on this question: (1) We establish that for grid-structured inputs not in general position, the separating capacity *l*\* equals the rank *R* of the input matrix. (2) We derive analytical formulae for the rank *R* of grid-like input matrices with integer periods, and generalize this notion to real-valued periods. (3) We show that the rank, and thus the separating capacity, asymptotically approaches the sum 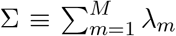 for generic real-valued periods. Our analytical results are verified by numerical simulation and counting (proofs provided in SI).

We begin with a numerical example for periods {3, 4}, Fig. 3A: The range over which all *K*-field arrangements are realizable – the place-cell separating capacity –is *l*\* = 6. Separating capacity curves for random inputs of the same input dimension (which equals matrix rank) as the grid-like input are also shown for comparison, Fig. 3B (solid and dashed gray curves, respectively). Although the separating capacity with grid-structured inputs is smaller than with inputs in general position (same input dimension), it is notably not much smaller, Fig. 3B (cf. black versus cyan) and it is larger than for dimension-matched inputs in general position if the readout weights are constrained to be non-negative (Fig. 3B, gray). (In a later section, we will show that the larger fraction of realizable field arrangements with random, general-position inputs are overall substantially less robust than with grid cell inputs.) Next, we show how to analytically characterize the separating capacity of place cells with grid-like inputs.

**Figure 3.**
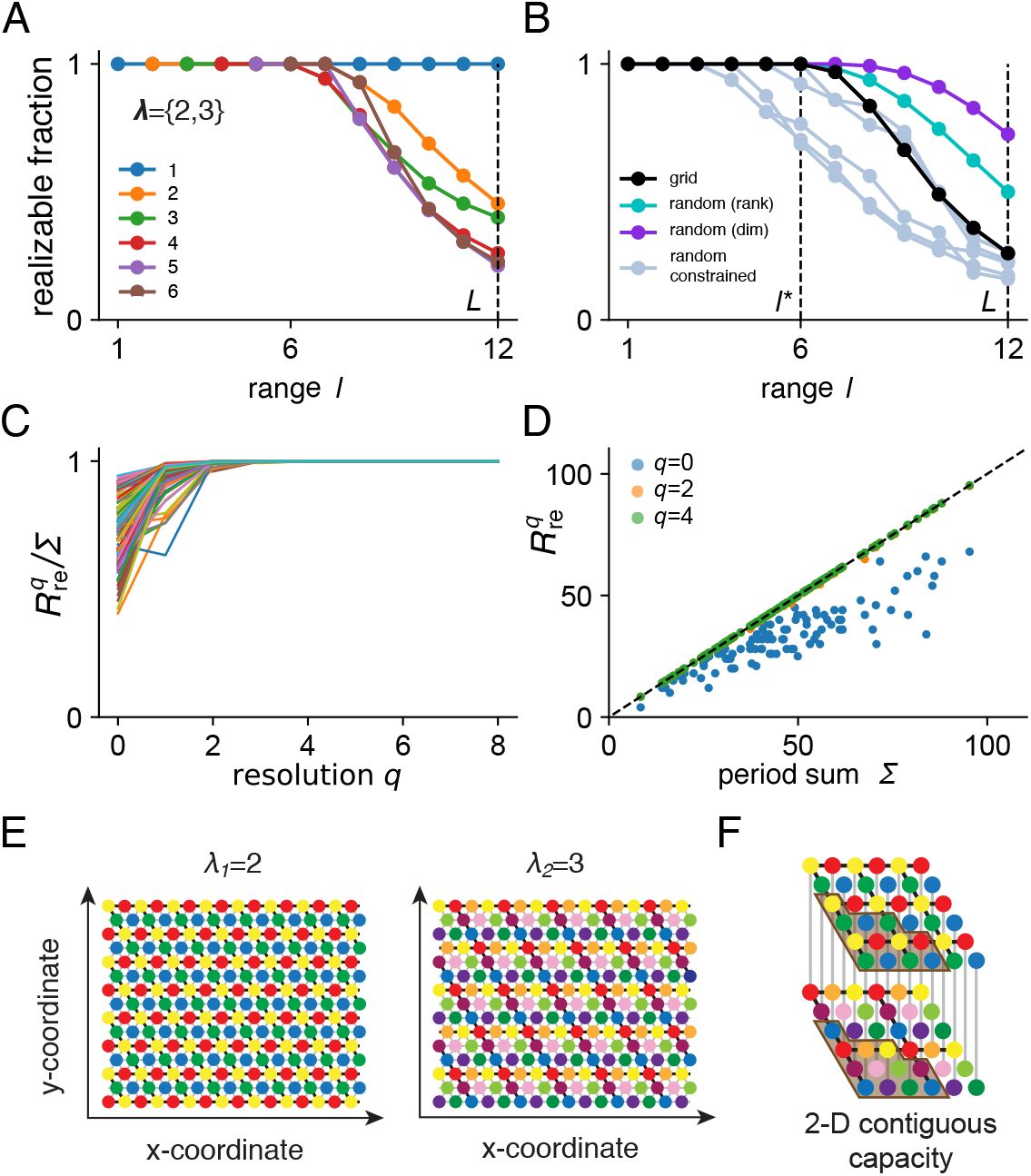
Place-cell separating capacity. (A) Fraction of realizable *K*-field arrangements vs track length (full range is marked *L*). Grid periods are {3, 4}. (B) Fraction of realizable field arrangements vs length of track (all *K*). Cyan (purple) line: fraction for random input of the same rank (dimension of input) as the grid-like input. Gray lines: fraction for random input of the same rank as the grid-like input with non-negative weight constraint. (C) Ratio of critical track length to sum of periods (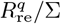, given by Eq. 6) as a function of number of decimal places *q* for 100 realizations of grid cell input with real-value periods, which are randomly drawn with the following constraints: *M* ∈ {2, 3, 4, 5, 6} and λ*_i_* ∈ [3, 20). The ratio approaches 1 from below as *q* increases. (D) Scatter plot of critical track length as a function of sum of grid periods. (E) The two spatial responses obtained for grid periods {2, 3}, which utilize 4 cells and 9 cells, respectively. (F) Maximal spatial mesh size for which each position has a unique grid-like code. The contiguous set of positions highlighted in brown achieves the separating capacity of the grid-like code.

### Separating capacity equals input-matrix rank

For inputs in general position, the separating capacity is equal to the rank of the input matrix (plus 1 when the threshold is allowed to be non-zero), which equals the dimension (number of cells) of the input patterns. When the inputs are not in general position, the separating capacity is upper-bounded by the rank, and the rank is upper-bounded by the dimension.

For any structured (non-general) input generated iteratively by repeated application of the same linear operator, the separating capacity saturates its bound and equals the input rank. To see this, consider a 1D translation-invariant code generated by successively applying a shift operator *J*, such that this process forms a sequence of distinct patterns ***x***, *J**x***, *J*^2^***x***, … *J^l^**x*** encoding *l* contiguous locations. The number of dimensions spanned by these patterns strictly increases upon adding a new pattern until some limit value *l*\*. After this, the dimension remains constant. This value equals the rank *R* of the input pattern matrix.

For a grid-like code, such an operator is given as a simultaneous one-unit phase shift in all grid modules. Thus, when *l* ≤ *R*, contiguous grid-like inputs are in general position and all dichotomies are realizable. Moreover, we show that when *l* = *R* + 1 input positions, there will be non-realizable field arrangements (SI Appendix). Therefore, the contiguous separating capacity is *l*\* = *R*.

Next, we compute this rank for the grid-like code under increasingly general assumptions.

### Matrix-rank formulae and convergence to sum of grid periods

#### Integer periods

We compute the input rank as the dimension of the vector space spanned by the rows of the input matrix. There are λ_*m*_ rows in the m-th module (1 ≤ *m* ≤ *M*), which consist of all the cyclic permutations of the same λ_*m*_-periodic binary vector of length *L* = LCM({λ_1_, …, λ_*M*_}). Thus, the subspaces generated by a module’s input rows have the convenient property to be stabilized by the circulant permutation matrix of size *L* × *L*. Therefore, the subspace spanned by each module can be represented as the span of a subset of those eigenvectors. By counting the number of unique basis elements generating all the subspaces using the inclusion-exclusion principle (SI Appendix), we get:

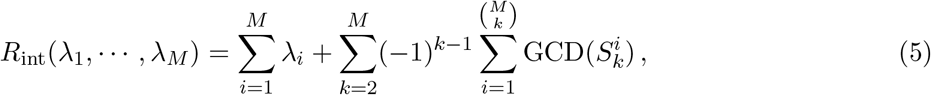

where 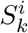 is the *i*-th *k*-element set of periods in {λ_1_, …, λ_*M*_}. It follows from the inclusion-exclusion principle that the rank satisfies 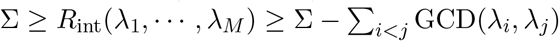. If the periods are pairwise-coprime, all the GCDs are 1 and this formula reduces to *R*_copr_(λ_1_, …, λ_*M*_) = Σ – *M* + 1. Thus, for large periods, *R*_copr_ approaches Σ, the sum of the grid-like input periods, which specifies the generic scaling of the rank. If selected uniformly at random over a finite range *N*, the module periods yield pairwise GCDs that scale as *o*(*N*) Cesaro (1881). Next, we exploit this fact to generalize the rank formula to real-valued periods. We will see that under reasonable conditions, the generalized rank and thus the separating capacity are well-approximated by Σ.

#### Real-valued periods

Actual grid periods are probably best described as truncated or finite-resolution versions of real-valued grid periods. We therefore consider the sequence of ranks 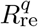 defined as:

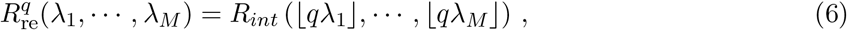

where ⌊·⌋ denotes the floor operation, *q* is an integer, and where we assume with no loss of generality that the periods are real numbers 0 < λ_1_ < … < λ_*M*_. The larger *q*, the finer the resolution of the integer approximation to the real-valued period, which intuitively corresponds to scaling the number of grid cells by *N* in each module. When the periods λ_1_ < … < λ_*M*_ are rational numbers, the sequence 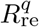 eventually scales linearly with *q*, when *q* exceeds their least common denominator. Thus, for rational periods, we can define the resolution-independent separating capacity as:

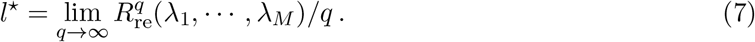

The above limit fails to exist for certain irrational numbers, thus extending our definition of separating capacity to real-valued periods must be done statistically. Assume with no loss of generality that the periods are drawn uniformly in (0, 1). Then, in the limit of fine resolution *q* → ∞, the truncations ⌊*q*λ_1_⌋, ⋯, ⌊*q*λ_*M*_⌋ are uniformly distributed integers in {1, …, *q*}, for which we have 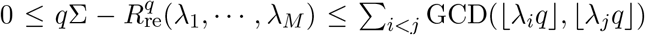. In the limit *q* → ∞, the probability that GCD(⌊λ_*i*_*q*⌋, ⌊λ_*j*_*q*⌋) = *g* asymptotically scales as 1/*g*^2^, which is independent of *q* Cesaro (1881). This implies that with probability one, the limit (7) is well-defined and amounts to *l*^⋆^ = Σ, which we equate with the resolution-independent separating capacity for real-valued periods. In Fig. 3C-D, we show numerically that this asymptotic scaling is actually reached quite rapidly with increasing *q*.

In short, all potential field configurations are realizable over a range given by the sum of periods of the grid modules contributing to that place cell’s response. Each local region of hippocampus likely receives inputs from two to three modules Honda et al. (2012); Cappaert et al. (2015), suggesting that the place-cell separating capacity is a few times the grid period.

The result above implies that place cells can only have complete flexibility in forming fields across one or at most a few small environments. Selecting an arrangement of fields over these spaces then constrains the choices that can be made over all remaining space. Crucially, if place cells maintain their input weights and thus a given arrangement for one environment, this choice restricts the field configurations in others, except for the fields driven directly by localized external sensory cues. Conversely, if a place cell learns field arrangements in a new environment by modifying the grid-to-place weights, it is highly likely that this learning will change the representation in previous environments (however, learning new associations between external cues and the place field scaffolds will not). Thus, the very small separating capacity of place cells according to our model may provide an explanation for the high volatility of the place code Ziv et al. (2013).

### Extension to 2D environments

The notion of contiguous separating capacity generalizes to more realistic spatial settings, including when the underlying space is 2-dimensional. In 2D, a grid cell’s peaks fall on a 2-dimensional triangular lattice, Fig. 3E. Our generalization is valid for d-dimensional (discretized) space. We previously defined the contiguous separating capacity as the maximum spatial distance (1D) over which all dichotomies of positions are linearly separable. More generally, we define the contiguous separating capacity as the volume of *d*-dimensional space over which all field arrangements are realizable with grid-like inputs, where each grid module is assumed to form a *d*-dimensional periodic response (using a number of cells that grows with *d* in the exponent), Fig. 3F. In dimension *d* > 1 the contiguous separating capacity can be achieved by several different sets of positions. For a set of grid modules with periods λ_1_, …, λ_*M*_ the separating capacity is 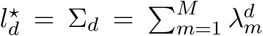, Fig. 3F (SI Appendix). This result follows from essentially the same reasoning as for 1D environments (SI Appendix), with the only added complication of considering the action of *d* commuting operators *J* on the grid-like code, with each operator associated with a one-unit shift along one of the spatial dimensions.

To summarize, the separating capacity of a place cell fed with grid cell input approaches the same capacity as if fed with general-position inputs of the same rank.

## Maximal margins and robustness

The margin of a decision boundary separating two classes of patterns is twice the shortest distance between the patterns and the boundary. The maximum margin is the largest achievable margin for that classification. The larger the maximum margin, the more robust is the desired classification to perturbations in the inputs or weights. For desired place-field arrangements, we thus compare maximum margins, herein simply referred to as margins, when the inputs are grid-like or not.

When the input patterns are structured rather than random, the margins take specific values based on the geometry of the input patterns rather than being described by distributions on statistical inputs (see e.g. Figure 2B). In general, the margin of a perceptron can be computed using a linear support vector machine (SVM)(Cortes and Vapnik, 1995), which reduces the problem to a quadratic programming problem. We use the SVC function in scikit-learn (Pedregosa et al., 2011) to numerically solve this problem for three types of input codes: the grid-like code, the shuffled grid-like code—a shuffled version of the grid-like code without modular structure–, and the random code of uniformly distributed random inputs. To make the comparison across codes fair, we generate: (1) patterns (columns) with the same total level of activity (their *L*1 norms sum to one and (2) the same total number of input patterns across codes (the number is smaller than the separating capacity for all three codes to ensure that a solution exists).

The random code admits more field arrangements (linearly separable dichotomies) with the same number of input patterns, as expected given that the random inputs are in general position, whereas the grid-like inputs are not (Fig. 4B). However, the grid-like code produces large margins compared to the random code, with substantially higher average and minimum maximum margins, Fig. 4A (cf. horizontal black lines and blue violins).

**Figure 4.**
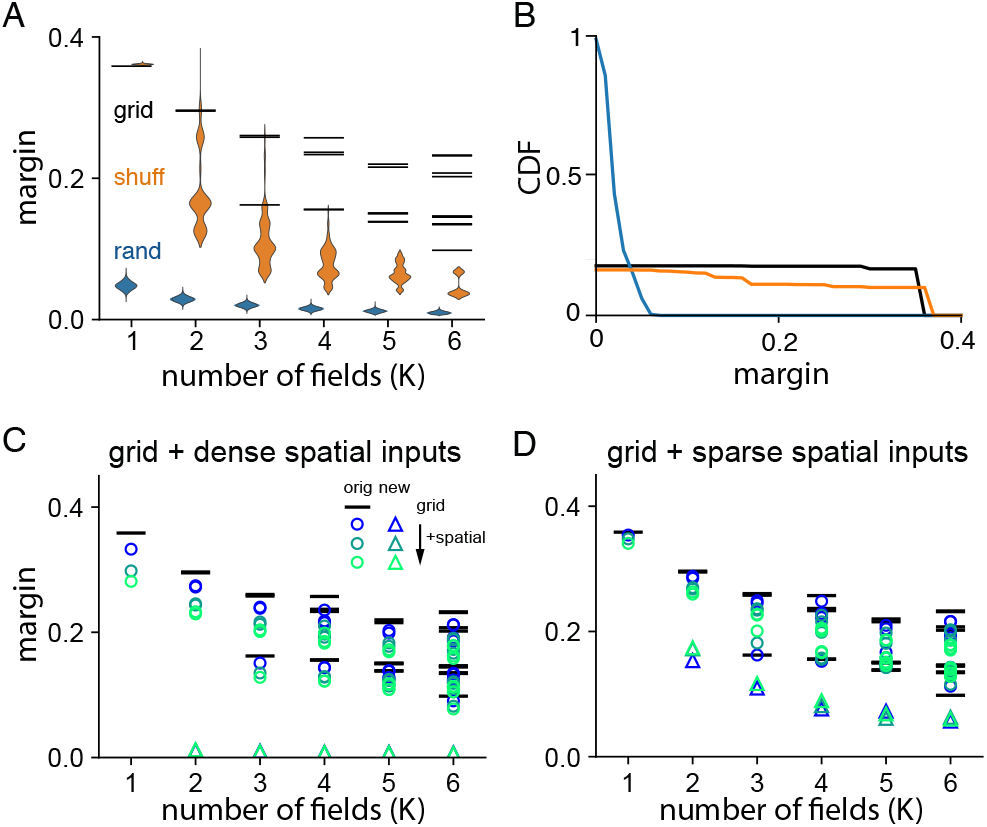
Maximum margins and robustness of place-field arrangements with grid and non-grid inputs. In all subplots, grid periods are {31, 43} and margins are determined using the SVM algorithm (with thresholds; no weight constraints). Input columns (patterns) are normalized to have unity *L*_1_ norm. (A) For grid-like inputs, the margins are deterministic and take up a small number of discrete values (black) across a very large number of field arrangements. Orange: Distribution of margins obtained when the grid-like input matrix is shuffled across columns (10 shuffles of the grid-like input matrix, sampling of 1000 realizable field arrangements per shuffle and per *K*). Blue: Distribution of margins for random inputs in general position (random input matrix sampled from a uniform distribution [0, 1); 10 realizations of a random matrix, 1000 samples per realization and *K*). (B) The cumulative distribution function (CDF) of margins for grid-like, shuffled, and random inputs. 1000 field arrangements are sampled for each *K* and the CDF includes margins across realizable field arrangements. Realizable field arrangements with grid-like inputs tend to have large margins; arrangements with shuffled, and random inputs in particular, have smaller margins. There are more field realizable arrangements with random inputs (fraction close to 1). (C) Effect of additional dense non-grid inputs on grid-like input margins: Black: margins as in (A-B). Blue, cyan, green circles: how the grid-like margins are modified when 30, 74, or 100 additional spatially dense inputs are provided. Each circle: average margin over 10 realizations of the input when separate dense spatial inputs are also included (dense spatial inputs drawn from a uniform distribution with ratio of peak random input to peak grid-like input 1:93). More field arrangements become realizable and the average margins over 10 randomly drawn newly realizable field arrangements are marked as triangles. (D) Same as (C) except that separate sparse spatial inputs are included (sparse spatial inputs of the same amplitude as the grid-like input are on at random locations with probability 1/190). Margins are determined largely by the grid cells.

The shuffled binary code exhibits a wider spread of margins (cf. orange violins), and thus overall smaller margins, than the grid-like code. In sum, the periodic and modular nature of the grid-like code creates wider margins and greater stability in the formation of place fields than alternative inputs.

Next we consider whether our result, that realizable place-fields occur in constrained arrangements, qualitatively persists with the addition of spatial inputs that are not grid-like. We added *L* non-grid spatially tuned inputs (with spatially dense tuning, modeled by frozen samples of uniformly distributed random inputs; *L* = 30, 74, 100, in addition to input from 74 grid cells, Fig. 4C. Alternatively, we add *L* spatially sparse non-grid inputs (same values for *L*; see Numerical Methods for details).

With either of these additional inputs, more field arrangements become realizable, as expected given that the input matrix is now of higher rank. With spatially dense inputs, the distribution of margins spreads, with a large number of small margins corresponding to the newly realizable field arrangements are substantially lower than the previous margins, Figs. 4C, so that the the number of robustly separable fields actually shrinks. With spatially sparse inputs, neither the number of arrangements nor their margins is strongly affected, meaning that the place-cell response remains largely determined by the grid cell drive.

Finally, real grid cells have graded responses profiles, with Gaussian-shaped activity bumps. These can be obtained by simple linear convolution of the discrete responses in our grid-like code with a smooth kernel. If the convolution matrix has rank equal to or larger than the binary grid cell input matrix, the set of realizable dichotomies remains unchanged.

In sum, random and shuffled input codes permit more field arrangements than grid-like inputs, but have smaller margins. The addition of a small random spatial input to the grid inputs also increases the number of field arrangements, but the additional achievable field arrangements have much smaller margins and are thus not robust (SI Appendix Fig. S6). The addition of non-grid spatial inputs, if dense, can increase the number of realizable arrangements, but reduce overall stability, while sparse non-grid spatial inputs do not significantly change the qualitative results with grid-like inputs alone.

## Predictions: Structure of the place-field scaffold

We have shown that place-field arrangements are highly constrained, such that only a tiny fraction of potential field arrangements within or across environments can be achieved. The ones that can be achieved can be understood intuitively with a simple picture: A field arrangement consists of the set of locations where a cell has fields, and equally important, the complementary set where it should have no fields. A place cell could choose its input weights and threshold to produce a field at one location, but that choice now strongly constrains other field and nonfield locations: Because grid-cell inputs are multiply peaked and non-local, strengthening weights from grid cells with certain phases and periods to obtain a peak at one location means that the place cell will also be strongly driven wherever subsets of those peaks recur in the grid input, as seen in Fig. 1D. This phenomenon, which we mathematically quantified above, generates predictions both about how place-field arrangements are constrained by their grid inputs, and about how place fields are constrained by others within a spatial scaffold. Overall, our model predicts that low-dimensional grid-cell states induce complex place cell activity patterns, rather than that place field patterns induce low-dimensional grid responses Dordek et al. (2016).

Our predictions are three-fold: (1) Place-field/grid-field relationships: The fields of a place cell will tend to coincide or align with the grid fields of the grid cells that drive it, within and across environments, Figure 5A. This means that all the fields of a place cell will fall at a field of at least one and likely more of the grid cells driving it, Fig. 5B. The stronger the input from a grid cell to a place cell, the more likely the place fields will coincide with that cell’s grid peaks. A corollary is that it should not be possible to induce stable place-field formation Bittner et al. (2015), except at locations consistent with the underlying grid cell input drive or where a strong local external cue is present. (2) Place field/place-field relationships, or the structure of the scaffold: The relative positions of the multiple fields of a place cell will be geometrically constrained, reflecting the combined geometries of the grid cells that project to it, with much more regularity than expected from random placement of fields. Specifically, the inter-field interval (IFI) distributions of the place fields in large environments will be structured with a few large components reflecting the combination of interfield intervals Yoon et al. (2016) in the underlying grid cells that drive the place cells, Figs. 5C-D. (3) More conceptually, our model predicts that external cues should recruit or “select” place fields based on the underlying grid-induced scaffold, because these fields are more robust, and also serve to associate external cues with internal motion-based estimates.

**Figure 5.**
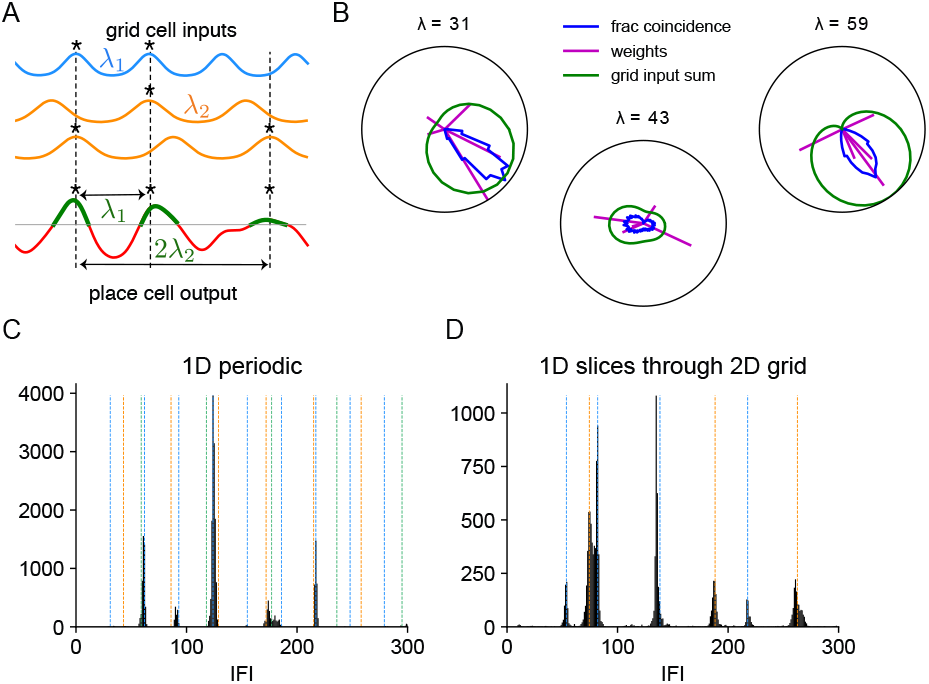
Predictions: Signatures of grid cell constraints on place-field arrangements. (A) Schematic: Our model, in which the majority of place fields are driven by grid cell inputs with Gaussian profile, makes predictions of two kinds, about place field-grid field relationships (vertical dashed lines) and place field-place field relationships (horizontal arrows). Place fields form when the summed input at a place cell is high (dashed lines). Thus, place fields tend to coincide with some activity peaks in some grid cells. (B) A place cell receives input from 15 grid cells from three modules with periods {31, 43, 59}, five in each module with randomly drawn phases. Weights from grid cells to the place cell are randomly drawn from Lognormal(0,1) (amplitudes relative to maximum weight in magenta), which determine the sum of periodic input each module contributes (with minimum of the sum subtracted and amplitudes relative to maximum in green). The polar coordinate represents phase *ϕ* ∈ [0, 1) and the radial coordinate displays the amplitude from 0 to 1. Place field locations are defined as the center of mass of the continuous pieces of input above the threshold, defined as mean plus two standard deviation of the total input across all locations. We compare the field locations with the firing fields of grid cells, defined as the locations above 0.95 of its peak amplitude. The blue line shows the fraction of coincident fields in a place cell with grid cells with all possible phases from three modules separately. Place fields in a place cell are likely to occur at field locations of a few subsets of grid cells with certain phases. (C-D) Predicted relationships between fields of a place cell. (C) Distribution of inter-field-intervals for adjacent fields in 100 model place cells with different weights as in (B). Peaks occur at integer combinations of the component grid periods (multiples of 31, 43, 59: blue, orange, green, respectively)(D) Same as (C) but the inputs correspond to 1D slices through 2D triangular grid-like responses with periods {31, 43}. Peaks are located at linear combinations of λ, 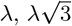 and 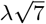 based on the spatial structure of grid cell activity along the slices.

## Discussion

### Summary of mathematical results

Mathematically, formulating the problem of place-field arrangements as a perceptron problem led us to examine the realizable (linearly separable) dichotomies of patterns that lie not in general position but on the vertices of regular polytopes, thus extending Cover’s results to a case of structured input Cover (1965). Input configurations not in general position complicate the counting of linearly separable dichotomies. For instance, counting the number of linearly separable Boolean functions, which is precisely the problem of counting the linearly separable dichotomies on the hypercube, is NP-hard Peled and Simeone (1985); Hegedus and Megiddo (1996).

For grid-like input, the regular polytope is obtained as an orthogonal product of simplices whose dimensions are set by the number of cells in each grid module. The grid-like input is a type of modular-one-hot code, in which the population is divided into modules and only one cell is active at a time per module. Exploiting the symmetries of modular-one-hot codes allowed us to characterize and enumerate the realizable *K*-dichotomies for small fixed cardinality *K* (*K* positive labels or *K*-field configurations), which for a fixed finite *K* is not a well-posed problem for patterns in general position since the solution will depend on the specific configuration of patterns. We characterized the realizable dichotomies by relying on combinatorial objects called Young diagrams Fulton and Fulton (1997). For the special case of *M* = 2 modules, we expressed the total number of dichotomies as a poly-Bernoulli number Kaneko (1997).

Interestingly, modular-one-hot codes allow each realizable dichotomy to equivalently be achieved with non-negative weights (SI Appendix), consistent with neurobiological constraints of excitatory projections from entorhinal cortex to hippocampus White et al. (1977); Cappaert et al. (2015).

### A large, rich scaffold of place field patterns for the formation of distinct maps

In our model, the dominant determinants of place field locations are grid cell inputs, under the reasoning that grid cells provide place cells with their primary source of spatial information based on integration of motion cues when external sensory cues are sparse. Below, we further examine this assumption. We showed that grid cell input enables the formation of a large number of place-field arrangements (Table 1). The number of realizable place-cell patterns when driven by grid cells far exceeds the number that can be generated recurrently purely within the hippocampus (assuming simple, non-modular recurrent dynamics within the hippocampus), as we argue below. Thus, the grid cell mechanism provides a quantifiably rich and large scaffold of distinct place-cell patterns, that are moreover robust, on which to “hang” external sensory cues and associate them with internal coordinates and each other in the formation of distinct maps for multiple environments.

As described in the section on separating capacity, once grid-to-place weights are set over a relatively small space, they set up a scaffold also outside of that space. Hanging an external cue would involve updating the weights from the external sensory inputs to place cells that are close to or above threshold based on the existing scaffold. This does not require relearning grid-to-place weights and does not cause interference with previously learned maps. By contrast, relearning the grid-to-place weights rearranges the overall scaffold, degrading previously learned maps (volatility Ziv et al. (2013)).

Note that all our results apply to the situation where grid cell states are incremented based on motion through cognitive spaces, not just physical space, and these states in turn drive place-cell responses (Killian et al., 2012; Constantinescu et al., 2016; Aronov et al., 2017).

### Other sources of spatial tuning in hippocampus

This work considers that place fields are essentially feedforward-driven conjunctions between (sparse) external sensory cues and (dense) motion-based internal position estimates represented periodically in grid cells. In considering place-cell responses as thresholded versions of their feedforward inputs, our model resembles notable other influential models in the literature, including Hartley et al. (2000); Solstad et al. (2006); Sreenivasan and Fiete (2011), as well as some contemporaneous models Whittington et al. (2019), and contrasts with other notable models in which place-cell responses emerge from recurrent pattern formation Tsodyks et al. (1996).

Our model is valid in the regime where grid cell inputs are the primary drivers of the spatially-tuned response of place cells. External sensory cues including environment or task boundaries and landmarks also provide reliable and spatially localized signals to drive place field firing (via lateral or medial entorhinal cortex); however, these external cues tend to be sparsely available, and we showed that sparse external cues do not qualitatively change our results and conclusions. Another possibility is that the hippocampus generates its own spatially tuned inputs through recurrent connections and internal velocity integration, according to the theory of pattern-forming continuous attractor networks Zhang (1996); Samsonovich and McNaughton (1997); Burak and Fiete (2009). However, the problem with these networks is that their dynamical capacity (number of fixed points) is very small (at most ~ *N* states with *N* neurons Amit et al. (1985); Gardner (1988); Abu-Mostafa and Jacques (1985); Sompolinsky and Kanter (1986); Samsonovich and McNaughton (1997); Battista and Monasson (2019)) unless they have special modular structure, as do grid cells, in which case the low-capacity individual modules together combinatorially produce a large library of states Burak and Fiete (2009); Sreenivasan and Fiete (2011); Mosheiff and Burak (2019); Fiete et al. (2014); Pogodin and Latham (2019); Chaudhuri and Fiete (2019). Thus, it is possible in principle that hippocampal cells could generate high-capacity stable spatial tuning without feedforward input from grid cells or external sensory cues if they contained modular organization, but without it, internal recurrent dynamics alone could not explain a large fraction of place fields.

In sum, we have shown that place cells can achieve a very large number of persistent and stable coding states by combining grid cell inputs, much larger than by general recurrent dynamics within the hippocampus, forming a large scaffold on which to associate external cues. Nevertheless, the allowed states are strongly constrained by the geometry of the grid-cell drive.

## Numerical methods

### Random input, weight-constrained random input and shuffled input

Entries of the random input matrix are uniformly distributed variables in [0,1). As far as linear separability is concerned (Fig. 3), matrices of the same rank and with the same number of patterns as the grid cell matrix are considered. As such input patterns are in general position, Cover’s counting theorem can be applied to count the realizable dichotomies Cover (1965). As for margins (Fig. 4), matrices of the same size as the grid cell matrix are used and threshold is imposed. Note that rank-constrained random input matrices simply result in smaller margins, which does not affect our conclusion that grid-like code gives rise to larger margins than random code. Weight-constrained random input (Figs. 3B,D) are random input with non-negative weight and threshold constraints imposed during training. As margins scale linearly with the linear spread of the patterns, input columns (patterns) are normalized to have unity *L*_1_ norm.

### Additional dense spatial inputs and sparse spatial inputs

On top of the grid cell input, additional spatial input is incorporated to test the robustness of the realizable field arrangements due to grid cell input (Figs. 4C,D). Dense spatial input consists of non-negative, uniformly distributed random variable with 20% of the mean activity of an average grid cell. Sparse spatial input consists of binary entries, with 20% of an average grid cell activity, randomly distributed within cells. All input columns (patterns) are normalized to have unity *L*_1_ norm.

### Gaussian-profiled grid cell input in a random weight model

The spatially periodic grid cell activity in 1D can be written as a function of phase (Fig. 5). The phase *ϕ* associated with location s and grid period λ is given by

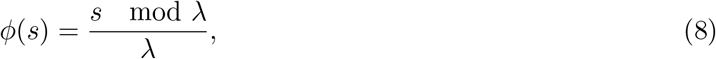

where 0 ≤ *ϕ* < 1. The activity profile of a grid cell is represented by a unimodal Gaussian in terms of phase (Sreenivasan and Fiete, 2011)

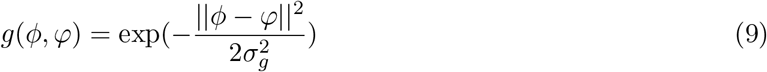

where

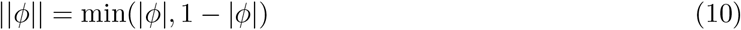

is the distance metric on phases. Each grid cell has an intrinsic phase *φ*. The width *σ_g_* in phase is independent of the underlying grid cell’s spatial response period and is set to be 0.16 (the full-width at half-max of the grid cell tuning curve equals 3/8 of the response period). Thus, the firing field width in real space scales linearly with the grid period. Extending to 2D, the activity of a grid cell at location **x** is given by

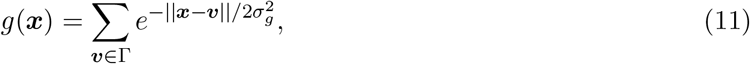

where 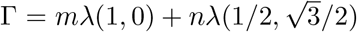 for all integers *m* and *n* Yoon et al. (2016). Concerning grid cell activity along a 1D slice through 2D with origin at *c* and orientation at *θ* relative to the *x*-axis, ***x*** can be parametrized by *t* as

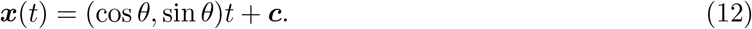

## Author contributions

I.R.F., T.T. and M.Y.Y. designed research; M.Y.Y. and T.T. performed research; M.Y.Y., L.A.S. and T.T. contributed new analytic tools; and T.T., M.Y.Y., and I.R.F. wrote the paper.

## Acknowledgements

This work was supported by the Simons Foundation through the Simons Collaboration on the Global Brain, the ONR, the Howard Hughes Medical Institute through the Faculty Scholars Program to IRF, and the Alfred P. Sloan Research Fellowship FG-2017-9554 to TT.

## Footnotes

1 Cover’s counting theorem is usually stated for perceptron with zero threshold for which the separating capacity is *N* rather than *N* + 1.

## References

Abu-Mostafa Y., and Jacques J.S. (1985). Information capacity of the Hopfield model. IEEE Transactions on Information Theory 31, 461–464.

Amit D.J., Gutfreund H., and Sompolinsky H. (1985). Storing Infinite Numbers of Patterns in a Spin-Glass Model of Neural Networks. Physical Review Letters 55, 1530–1533.

Amit D.J., Wong K.Y.M., and Campbell C. (1989). Perceptron learning with sign-constrained weights. Journal of Physics A: Mathematical and General 22, 2039–2045.

Aronov D., Nevers R., and Tank D.W. (2017). Mapping of a non-spatial dimension by the hippocampal-entorhinal circuit. Nature 543, 719–722.

Battista A., and Monasson R. (2019). Capacity-resolution trade-off in the optimal learning of multiple low-dimensional manifolds by attractor neural networks.

Bittner K.C., Grienberger C., Vaidya S.P., Milstein A.D., Macklin J.J., Suh J., Tonegawa S., and Magee J.C. (2015). Conjunctive input processing drives feature selectivity in hippocampal CA1 neurons. Nature Neuroscience 18, 1133–1142.

Brunel N., Hakim V., Isope P., Nadal J.P., and Barbour B. (2004). Optimal Information Storage and the Distribution of Synaptic Weights: Perceptron versus Purkinje Cell. Neuron 43, 745–757.

Burak Y., and Fiete I.R. (2009). Accurate Path Integration in Continuous Attractor Network Models of Grid Cells. PLoS Computational Biology 5, e1000291.

Burgess N. (2008). Grid cells and theta as oscillatory interference: Theory and predictions. Hippocampus 18, 1157–1174.

Cadena C., Carlone L., Carrillo H., Latif Y., Scaramuzza D., Neira J., Reid I., and Leonard J.J. (2016). Past, present, and future of simultaneous localization and mapping: Toward the robust-perception age. Trans. Rob. 32, 1309–1332.

Cappaert N.L.M., Van Strien N.M., and Witter M.P. (2015). Chapter 20 - Hippocampal Formation (San Diego: Academic Press), pp. 511–573.

Cesaro E. (1881). Démonstration élémentaire et généralisation de quelques théoremes de M. Berger. Mathesis 1, 99–102.

Chaudhuri R., and Fiete I. (2019). Bipartite expander Hopfield networks as self-decoding high-capacity error correcting codes. In Advances in Neural Information Processing Systems 32, H. Wallach, H. Larochelle, A. Beygelzimer, F. d\textquotesingle Alche-Buc, E. Fox, and R. Garnett, eds. (Curran Associates, Inc.), pp. 7686–7697.

Cheung A., Ball D., Milford M., Wyeth G., and Wiles J. (2012). Maintaining a Cognitive Map in Darkness: The Need to Fuse Boundary Knowledge with Path Integration. PLOS Computational Biology 8, e1002651.

Colgin L.L., Moser E.I., and Moser M.B. (2008). Understanding memory through hippocampal remapping. Trends Neurosci 31, 469–77.

Constantinescu A.O., O’Reilly J.X., and Behrens T.E.J. (2016). Organizing conceptual knowledge in humans with a gridlike code. Science 352, 1464 LP – 1468.

Cortes C., and Vapnik V. (1995). Support-Vector Networks. Machine Learning 20, 273–297.

Cover T.M. (1965). Geometrical and Statistical Properties of Systems of Linear Inequalities with Applications in Pattern Recognition. IEEE Transactions on Electronic Computers EC-14, 326–334.

de Andrade R.F., Lundberg E., and Nagle B. (2015). Asymptotics of the extremal excedance set statistic. European Journal of Combinatorics 46, 75–88.

Dordek Y., Soudry D., Meir R., and Derdikman D. (2016). Extracting grid cell characteristics from place cell inputs using non-negative principal component analysis. eLife 5, e10094.

Fenton A.A., Kao H.Y., Neymotin S.A., Olypher A., Vayntrub Y., Lytton W.W., and Ludvig N. (2008). Unmasking the CA1 Ensemble Place Code by Exposures to Small and Large Environments: More Place Cells and Multiple, Irregularly Arranged, and Expanded Place Fields in the Larger Space. The Journal of Neuroscience 28, 11250.

Fiete I., Schwab D.J., and Tran N.M. (2014). A binary hopfield network with 1/log(n) information rate and applications to grid cell decoding.

Fiete I.R., Burak Y., and Brookings T. (2008). What Grid Cells Convey about Rat Location. Journal of Neuroscience 28, 6858–6871.

Fulton W., and Fulton M.W. (1997). Young tableaux: with applications to representation theory and geometry, vol. 35

Gardner E. (1988). The space of interactions in neural network models. Journal of Physics A: Mathematical and General 21, 257–270.

Hafting T., Fyhn M., Molden S., Moser M.B., and Moser E.I. (2005). Microstructure of a spatial map in the entorhinal cortex. Nature 436, 801–806.

Hartley T., Burgess N., Lever C., Cacucci F., and O’Keefe J. (2000). Modeling place fields in terms of the cortical inputs to the hippocampus. Hippocampus 10, 369–379.

Hegedus T., and Megiddo N. (1996). On the geometric separability of boolean functions. Discrete Applied Mathematics 66, 205–218.

Hollup S.A., Molden S., Donnett J.G., Moser M.B., and Moser E.I. (2001). Accumulation of hippocampal place fields at the goal location in an annular watermaze task. The Journal of neuroscience: the official journal of the Society for Neuroscience 21, 1635–44.

Honda Y., Sasaki H., Umitsu Y., and Ishizuka N. (2012). Zonal distribution of perforant path cells in layer III of the entorhinal area projecting to CA1 and subiculum in the rat. Neuroscience Research 74, 200–209.

Itskov V., and Abbott L.F. (2008). Pattern Capacity of a Perceptron for Sparse Discrimination. Physical Review Letters 101, 18101.

Kaneko M. (1997). Poly-bernoulli numbers. Journal de theorie des nombres de Bordeaux 9, 221–228.

Kanitscheider I., and Fiete I. (2017a). Emergence of dynamically reconfigurable hippocampal responses by learning to perform probabilistic spatial reasoning. bioRxiv p. 231159.

Kanitscheider I., and Fiete I. (2017b). Making our way through the world: Towards a functional understanding of the brain’s spatial circuits. Current Opinion in Systems Biology 3, 186–194.

Kanitscheider I., and Fiete I. (2017c). Training recurrent networks to generate hypotheses about how the brain solves hard navigation problems. Advances in Neural Information Processing Systems pp. 4529–4538.

Killian N.J., Jutras M.J., and Buffalo E.A. (2012). A map of visual space in the primate entorhinal cortex. Nature 491, 761–4.

Leonard J.J., and Durrant-Whyte H.F. (1991). Mobile robot localization by tracking geometric beacons. IEEE Transactions on Robotics and Automation 7, 376–382.

Mathis A., Herz A.V.M., and Stemmler M. (2012). Optimal Population Codes for Space: Grid Cells Outperform Place Cells. Neural Computation 24, 2280–2317.

McNaughton B.L., Battaglia F.P., Jensen O., Moser E.I., and Moser M.B. (2006). Path integration and the neural basis of the ‘cognitive map’. Nature Reviews Neuroscience 7, 663–678.

Milford M.J., Wyeth G.F., and Prasser D. (2004). RatSLAM: a hippocampal model for simultaneous localization and mapping. In IEEE International Conference on Robotics and Automation, 2004. Proceedings. ICRA ’04. 2004. vol. 1, pp. 403–408 Vol.1.

Mosheiff N., and Burak Y. (2019). Velocity coupling of grid cell modules enables stable embedding of a low dimensional variable in a high dimensional neural attractor. eLife 8, e48494.

O’Keefe J. (1976). Place units in the hippocampus of the freely moving rat. Experimental Neurology 51, 78–109.

O’Keefe J., and Burgess N. (1996). Geometric determinants of the place fields of hippocampal neurons. Nature 381, 425–428.

O’Keefe J., and Dostrovsky J. (1971). The hippocampus as a spatial map. Preliminary evidence from unit activity in the freely-moving rat. Brain research 34, 171–5.

O’Keefe J., and Nadel L. (1978). The hippocampus as a cognitive map (Clarendon Press).

Park E., Dvorak D., and Fenton A.A. (2011). Ensemble Place Codes in Hippocampus: CA1, CA3, and Dentate Gyrus Place Cells Have Multiple Place Fields in Large Environments. PLoS ONE 6, e22349.

Pedregosa F., Varoquaux G., Gramfort A., Michel V., Thirion B., Grisel O., Blondel M., Prettenhofer P., Weiss R., Dubourg V., Vanderplas J., Passos A., Cournapeau D., Brucher M., Perrot M., and Duchesnay E. (2011). Scikit-learn: Machine learning in Python. Journal of Machine Learning Research 12, 2825–2830.

Peled U.N., and Simeone B. (1985). Polynomial-time algorithms for regular set-covering and threshold synthesis. Discrete Applied Mathematics 12, 57 – 69.

Pogodin R., and Latham P. (2019). Memories in coupled winner-take-all networks. In Cosyne Abstracts.

Postnikov A. (2006). Total positivity, grassmannians, and networks. arXiv preprint math/0609764

Quirk G.J., Muller R.U., and Kubie J.L. (1990). The firing of hippocampal place cells in the dark depends on the rat&#039;s recent experience. The Journal of Neuroscience 10, 2008 LP – 2017.

Rich P.D., Liaw H.P., and Lee A.K. (2014). Large environments reveal the statistical structure governing hippocampal representations. Science 345.

Rosenblatt F. (1958). The perceptron: a probabilistic model for information storage and organization in the brain. Psychological review 65, 386–408.

Samsonovich A., and McNaughton B.L. (1997). Path integration and cognitive mapping in a continuous attractor neural network model. The Journal of neuroscience: the official journal of the Society for Neuroscience 17, 5900–20.

Solstad T., Moser E.I., and Einevoll G.T. (2006). From grid cells to place cells: A mathematical model. Hippocampus 16, 1026–1031.

Sompolinsky H., and Kanter I. (1986). Temporal Association in Asymmetric Neural Networks. Physical Review Letters 57, 2861–2864.

Sreenivasan S., and Fiete I. (2011). Grid cells generate an analog error-correcting code for singularly precise neural computation. Nature Neuroscience 14, 1330–1337.

Tolman E.C. (1948). Cognitive maps in rats and men. Psychological Review 55.

Tsodyks M.V., Skaggs W.E., Sejnowski T.J., and McNaughton B.L. (1996). Population dynamics and theta rhythm phase precession of hippocampal place cell firing: a spiking neuron model. Hippocampus 6, 271–80.

Vapnik V.N. (1998). Statistical learning theory (Wiley).

Welinder P.E., Burak Y., and Fiete I.R. (2008). Grid cells: The position code, neural network models of activity, and the problem of learning. Hippocampus 18, 1283–1300.

White W.F., Nadler J.V., Hamberger A., Cotman C.W., and Cummins J.T. (1977). Glutamate as transmitter of hippocampal perforant path. Nature 270, 356–357.

Whittington J.C.R., Muller T.H., Mark S., Chen G., Barry C., Burgess N., and Behrens T.E.J. (2019). The Tolman-Eichenbaum Machine: Unifying space and relational memory through generalisation in the hippocampal formation. bioRxiv p. 770495.

Widloski J., and Fiete I. (2014). How Does the Brain Solve the Computational Problems of Spatial Navigation? BT - Space, Time and Memory in the Hippocampal Formation (Springer Vienna), pp. 373–407.

Yim M.Y., Taillefumier T., and Fiete I.R. (2019a). A robust signature of grid code readout in place field statistics. In Conference on Learning and Memory at UT Austin abstract.

Yim M.Y., Taillefumier T., and Fiete I.R. (2019b). Mechanistic models of place cell statistics in large environments. In SfN abstract.

Yoon K., Buice M.A., Barry C., Hayman R., Burgess N., and Fiete I.R. (2013). Specific evidence of low-dimensional continuous attractor dynamics in grid cells. Nature Neuroscience 16, 1077–1084.

Yoon K., Lewallen S., Kinkhabwala A.A., Tank D.W., and Fiete I.R. (2016). Grid Cell Responses in 1D Environments Assessed as Slices through a 2D Lattice. Neuron 89, 1086–1099.

Zhang K. (1996). Representation of spatial orientation by the intrinsic dynamics of the head-direction cell ensemble: a theory. The Journal of neuroscience: the official journal of the Society for Neuroscience 16, 2112–26.

Ziv Y., Burns L.D., Cocker E.D., Hamel E.O., Ghosh K.K., Kitch L.J., Gamal A.E., and Schnitzer M.J. (2013). Long-term dynamics of CA1 hippocampal place codes. Nature Neuroscience 16, 264–266.

Zuev Y.A. (1989). Asymptotics of the logarithm of the number of threshold functions of the algebra of logic, vol. 39.

